# Identification of kinase inhibitors as potential host-directed therapies for intracellular bacteria

**DOI:** 10.1101/2023.08.28.555045

**Authors:** Robin H.G.A. van den Biggelaar, Kimberley V. Walburg, Susan van den Eeden, Cassandra L.R. van Doorn, Eugenia Meiler, Alex S. de Ries, M. Chiara Fusco, Annemarie H. Meijer, Tom H.M. Ottenhoff, Anno Saris

## Abstract

The emergence of antimicrobial resistance has created an urgent need for alternative treatment strategies against deadly bacterial species. In this study, we investigated the potential of kinase inhibitors as host-directed therapies (HDTs) for combating infectious diseases caused by intracellular bacteria, specifically *Salmonella* Typhimurium (*Stm*) and *Mycobacterium tuberculosis* (*Mtb*). We screened 827 ATP-competitive kinase inhibitors with known target profiles from two Published Kinase Inhibitor Sets (PKIS1 and PKIS2) using intracellular infection models for *Stm* and *Mtb*, based on human cell lines and primary macrophages. Additionally, the *in vivo* efficacy of the compounds was assessed using zebrafish embryo infection models. Our kinase inhibitor screen identified 14 hit compounds for *Stm* and 19 hit compounds for *Mtb* that were effective against intracellular bacteria and non-toxic for host cells. Further validation experiments showed the high efficacy of most *Stm* hit compounds and their ability to fully clear the intracellular infection both in cell lines and primary human macrophages. From these, two structurally related *Stm* hit compounds, GSK1379738A and GSK1379760A, exhibited significant effectiveness against *Stm* in infected zebrafish embryos. Compounds that were active against intracellular *Mtb* included morpholino-imidazo/triazolo-pyrimidinones that specifically target the kinases PIK3CB and PIK3CD as well as 2-aminobenzimidazoles targeting BLK, ABL1 and TRKA. Overall, this study provided insight into critical kinase targets acting at the host-pathogen interface and identified novel kinase inhibitors as potential HDTs for intracellular bacterial infections.

## Introduction

Infectious diseases caused by intracellular bacteria such as *Salmonella* species and *Mycobacterium tuberculosis* (*Mtb*) significantly impact global health. *Salmonella* serotypes are categorized into two main groups based on their clinical symptoms: (para)typhoid and non-typhoid, which are associated with (para)typhoid fever and gastroenteritis, respectively. These pathogens result in over 100 million cases of *Salmonella*-related diseases worldwide, with approximately 250,000 deaths annually [1,2]. *Mtb* is estimated to be latently present in almost a quarter of the world’s population, with around 10 million new cases of tuberculosis (TB) each year, and around 1.5 million deaths [2–4]. Despite a previous decline in TB cases, the COVID-19 pandemic has had an impact on the management of TB diagnosis and treatment, resulting in a recent increase in TB cases [4]. Another significant challenge in both *Salmonella* and *Mtb* infections is the rise of antimicrobial resistance, which highlights the need for alternative therapeutic approaches [1,4,5].

As both *Salmonella* species and *Mtb* are intracellular pathogens that depend on host cells for survival and replication, host-directed therapy (HDT) may be a promising avenue to treat these infections. HDT aims to target the host response against infection rather than the pathogen itself and may be used as an adjunctive or alternative treatment to classical antibiotics. Importantly, HDTs have the potential to target antibiotic resistant strains. Due to their involvement in many host-pathogen interaction mechanisms, kinase inhibitors have previously gained attention as HDT candidates against intracellular bacteria. Both *Salmonella enterica* serovar Typhimurium (*Stm*) and *Mtb* activate the PI3K/Akt signaling pathway through their virulence factors to stimulate host cell survival. Consequently, chemical inhibition (e.g. H-89 and AR-12) or genetic inhibition of kinases that are part of this signaling pathway impairs bacterial growth [6–8]. Furthermore, chemical or genetic inhibition of several receptor tyrosine kinases stimulates host cell control of intracellular *Mtb* [9]. For instance, inhibition of ABL1 by imatinib consistently reduces intracellular *Mtb* burden, most likely by overcoming pathogen-driven suppression of lysosomal acidification [6,9–13]. Currently, ABL1 inhibitor imatinib is being tested in clinical trials as adjunctive therapy together with an antibiotic regimen of rifabutin and isoniazid against drug-sensitive TB [14].

In this study, we explored the potential of very well characterized kinase inhibitors as candidates for HDT by screening 827 ATP-competitive kinase inhibitors of two Published Kinase Inhibitor Sets, PKIS1 and PKIS2, in intracellular infection models for *Stm* and *Mtb* [15,16]. Moreover, we evaluated the most promising compounds *in vivo* in zebrafish embryo models. The findings of this study resulted in the identification of novel HDTs and well as HDT targets critical for host-pathogen-interactions that will help to overcome the challenges posed by emerging antibiotic resistance.

## Materials and Methods

### Ethical statements

Human blood samples were isolated from buffy coats obtained from healthy donors after written informed consent (Sanquin, Amsterdam, the Netherlands). The biological samples were sourced ethically and their research use was in accordance with the terms of the informed consents under an IRB/EC approved protocol. The husbandry of adult zebrafish lines described in this study was in accordance with guidelines from the local animal welfare committee (DEC) of Leiden University (License number: protocol 14,198), and in compliance with the international guidelines specified by the EU Animal Protection Directive 2010/63/EU. Experiments with zebrafish embryos were performed within 5 days post fertilization and therefore did not involve any procedures within the meaning of Article 3 of Directive 2010/63/EU.

### Reagents

Kinase inhibitor H-89 dihydrochloride was purchased from Sigma-Aldrich (Merck, Darmstadt, Germany). The H-89 analogue 97i was a gift from prof. dr. M. van der Stelt and prepared *in house* according to a previous publication [17,18]. Rifampicin, moxifloxacin hydrochloride, gentamycin sulfate, dimethyl sulfoxide (DMSO) and Triton X-100 were all purchased from Sigma-Aldrich. A total of 827 kinase inhibitors from the published kinase inhibitor sets (PKIS)1 [15] and PKIS2 [16] was obtained from GlaxoSmithKline (GSK) and the Structural Genomics Consortium of the University of North Carolina at Chapel Hill (SGC-UNC). All kinase inhibitors, including H-89 and 97i, were dissolved at 10 mM concentration in DMSO. Mouse anti-human antibodies CD14-FITC (clone 63D3), CD163-PE (clone GHI/61), CD14-PE/Cy7 (clone 63D3) and CD1a-AF647 (clone HI149) were purchased from Biolegend (San Diego, CA, USA). Mouse anti-human CD11b-BB515 (clone ICRF44) was purchased from BD Biosciences (Franklin Lakes, NJ, USA).

### Cell culture

HeLa epithelial cells and MelJuSo human melanoma cells were cultured in Gibco Iscove’s Modified Dulbecco’s Medium (IMDM; ThermoFisher Scientific, the Netherlands) supplemented with 10% fetal bovine serum (FBS; Greiner Bio-One, Alphen a/d Rijn, the Netherlands), 100 units/ml Gibco penicillin and 100 µg/ml Gibco streptomycin (both from ThermoFisher Scientific) at 37°C/5% CO_2_.

Primary human macrophages were generated by differentiating blood-derived CD14^+^ monocytes for 6 days in Roswell Park Memorial Institute (RPMI)-1640 medium (ThermoFisher Scientific) containing 10% HyClone FBS (Cytiva, MA, Marlborough, USA), 100 units/ml penicillin, 100 µg/ml streptomycin and either 5 ng/ml granulocyte-macrophage colony-stimulating factor (GM-CSF; R&D Systems, Abingdon, UK) to promote M1-differentiation or 20 ng/ml macrophage colony-stimulating factor (M-CSF; R&D Systems) to promote M2-differentiation as previously described [9]. Macrophages were collected by incubation in Gibco Trypsin-EDTA solution (ThermoFisher Scientific) and gentle scraping. The M1 and M2 macrophage phenotypes were validated based on morphology and surface marker expression using flow cytometry with M1 macrophages being CD14^low^, CD163^low^ and CD11b^high^, whereas M2 macrophages were CD14^high^, CD163^high^, CD11b^low^ [19].

### Bacterial culture

*Stm* strain SL1344 with plasmid pMW211[C.10E/DsRed] [20] and strain 12023 with plasmid pluxCDABE [6] were recovered from frozen glycerol stock and cultured in Difco Luria-Bertani (LB) Broth (BD Biosciences) containing 100 µg/ml ampicillin (Merck, Darmstadt, Germany) overnight at 37°C in a shaking incubator. *Stm* was subcultured 1:33 three to four hours prior to infection to obtain a log-phase bacterial culture. *Mtb* strain H37Rv with plasmid pSMT3[Phsp60/DsRed] [9] was cultured at 37°C in complete Difco Middlebrook 7H9 Broth (BD Biosciences), supplemented with 10% ADC (BD Biosciences), 0.5% Tween-80 (Sigma-Aldrich), 2% glycerol (Sigma-Aldrich) and 50 µg/ml Gibco hygromycin (ThermoFisher Scientific) in a shaking incubator. *Mtb* was split once a week to maintain log-phase bacterial cultures. *Mycobacterium marinum* (*Mmar*) strain M with plasmid pTEC15[mWasabi] [21] was diluted to an OD_600_ of 0.1 the day before infection and cultured overnight complete 7H9 medium at 28.5°C in a static incubator.

### *Stm* and *Mtb* intracellular infection and HDT treatment

HeLa or MelJuSo cells were resuspended in IMDM with 10% FBS without antibiotics and seeded with 10,000 cells/well and 9000 cells/well, respectively, into Costar flat-bottom 96-well plates (Corning, Amsterdam, the Netherlands). M1 and M2 macrophages were resuspended in RPMI with 10% FBS without antibiotics and seeded with 30,000 cells/well into Costar flat-bottom 96-well plates. The cells were incubated overnight at 37°C/5% CO_2_. The next day, cells were inoculated with log-phase bacterial suspensions in cell culture medium at an expected multiplicity of infection (MOI) of 10. Accuracy of the MOI was validated by plating serial dilutions of the *Stm* and *Mtb* inoculums on Difco LB agar plates and Middlebrook 7H10 agar plates supplemented with 10% OADC and 5% glycerol, respectively (all from BD Biosciences). The observed MOIs were 11.6 (n = 18; range 3.6 – 15.9) and 18.9 (n = 13; range 5.2 – 41.1), respectively. Plates with infected cells were centrifuged for 3 min at 150 *g* and incubated for 20 min for *Stm* infection or 1 hour for *Mtb* infection at 37°C/5% CO_2_. Extracellular bacteria were removed by incubation with fresh cell culture medium supplemented with 30 µg/ml gentamicin sulphate (Lonza BioWhittaker, Basel, Switzerland) for 15 min at 37°C/5% CO_2_. Infected cells were incubated overnight at 37°C/5% CO_2_ in cell culture medium in the presence of kinase inhibitors at 10 µM concentration and 5 µg/ml gentamycin sulphate-containing medium to prevent growth of extracellular bacteria. Kinase inhibitors H-89 and its derivative 97i were used at 10 µM concentrations as HDT positive controls for *Stm* and *Mtb*, respectively, and an equal amount of DMSO (% v/v), used as solvent for all compounds, was used as a negative control. Rifampicin and gentamycin were used at 1 µM and 30 µg/ml, respectively.

### Compound screens by flow cytometry

HeLa and MelJuSo cells were harvested by trypsinization and fixed with 1% paraformaldehyde. Samples from the primary screen were measured on a FACSCalibur with high-throughput samples (HTS) extension and samples from the rescreen were measured on a FACSLyric (BD Biosciences) with HTS extension (all from BD Biosciences). Flow cytometry data was analysed using FlowJo version 10 (TreeStar, Ashland, OR, USA). Each plate contained H-89 and DMSO as positive and negative controls, and each plate was tested in triplicate. In addition, the rescreens included H89-derivative 97i as a more suitable positive control for intracellular *Mtb*, since 97i was recently shown to be more effective than H89 [18]. *Mtb*-infected MelJuSo cells positive for DsRed were gated to calculate the percentage of infected cells. For *Stm*-infected HeLa cells, a DsRed-bright population could be distinguished from a DsRed-dim population (**Supplementary Figure 1a**). Both the total *Stm*-DsRed+population and the DsRed-bright population were initially used as readouts for the primary screen, but eventually the DsRed-bright population was used for selection of hit compounds since this population was found to contain nearly all intracellular bacteria after FACS sorting (see methods below; **Supplementary Figure 2**). The cell count was used as readout of cell viability.

### Data analysis of flow cytometry screens

Standard z-scores were calculated from the flow cytometry data to identify candidate ‘hit’ compounds by z = (x-μ)/σ – z_neg_, where x is total event count or the percent of DsRed-bright cells or DsRed+ cells from a single well, μ is the mean from wells of the plate, σ is the standard deviation of the plate and z_neg_ is the mean z-score of the negative control of the plate (i.e. DMSO). The subtraction of z_neg_ was used to correct z-scores for plate-to-plate variations in baseline infections rates or cell viability. Values that deviated two-fold from the plate mean were excluded to calculate the plate mean and standard deviation as used in the aforementioned formula, to minimize the possibility that the presence of extreme values on a plate affected the plate mean and standard deviation compared to the other plates. Compounds that had a Z-score < –3 for cell count were considered cytotoxic and were therefore excluded as hit compounds. Compounds with a z-score below or above the critical value of respectively –2 or 2 for DsRed+ and/or DsRed-bright events and a z-score above the critical value of –3 for cell count were considered ‘hit’ compounds. However, after the rescreen only compounds with a DsRed z-score below the critical value of –2 were further analysed.

### Identification of kinase targets

Kinase inhibition data of PKIS compounds at 1 µM concentration has previously been determined in biochemical assays by others [15,16]. KinMap phylogenetic trees of protein kinase families were generated at https://kinhub.org/kinmap/ for visual representation of kinase inhibition data [22]. A phylogenetic tree of phosphatidylinositol kinases was created by multiple sequence alignment using Clustal Omega at https://www.ebi.ac.uk/Tools/msa/clustalo/, based on the PROSITE-predicted kinase catalytic domains, and neighbor-joining at https://icytree.org/ [23]. Most PKIS compounds target multiple kinases. The contribution of individual kinases to host-pathogen interactions was determined by genetic kinase inhibition data from a previous publication in which siKinome screens were performed on the HeLa-*Stm* and MelJuSo-*Mtb* intracellular infection models [9].

### Colony-forming unit assay

Infected cells were lysed in sterile water + 0.05% Invitrogen UltraPure SDS solution (ThermoFisher Scientific) to release bacteria. Bacterial suspensions and lysates of cells infected with *Stm* and *Mtb* were 5-fold serially diluted and 10-μl drops were plated on square agar plates, using Difco LB agar plates for *Stm* and Middlebrook 7H10 agar plates for *Mtb*. The plates were left to dry and incubated overnight at 37°C/5% CO_2_ for *Stm* and approximately two weeks for *Mtb*, after which colony forming units (CFUs) were counted manually.

### FACS sorting

*Stm*-infected HeLa cells were FACS-sorted in DsRed-dim and DsRed populations using the CytoFLEX SRT II cell sorter (Beckman Coulter Fullerton, CA, USA) of the Flow Cytometry Core Facility of the Leiden University Medical Centre to determine the intracellular bacterial burden of both populations. After sorting, the cells were lysed and the lysate was plated out on LB agar plates to determine the CFU count as described above.

### Bacterial growth assay

*Stm* or *Mtb* cultures at a concentration corresponding to an absorbance of 0.1 at 600 nm wavelength were incubated with the kinase inhibitors at 10 μM in flat-bottom 96-well plates. The absorbance was measured using an EnVision plate reader (PerkinElmer, Waltham, MA, USA). The plates were incubated at 37 °C overnight for *Stm* and for a period of 15 days for *Mtb*. The absorbance was measured again the next day for *Stm* and every two days for *Mtb*.

### Lactate dehydrogenase release cytotoxicity assay

Before harvesting or lysing cells of intracellular infection assays, supernatant from the cells was collected and used to quantify LDH release using the LDH cytotoxicity detection kit (Roche, Merck, Darmstadt, Germany) according to the manufacturer’s instructions. Quantification was performed with an EnVision plate reader or SpectraMax i3x (Molecular Devices, San Jose, CA, USA) plate reader by using the absorbance at 485 nm as signal and the absorbance at 690 nm as reference wavelength for background subtraction. The DMSO solvent control was used as negative control. Triton X-100 results in maximum LDH release by permeabilizing the cells and was used as positive control. The cell viability was calculated using the following formula: 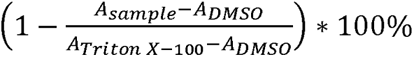 with *A* for absorbance at 485 nm after subtraction of the absorbance at 690 nm.

### Bioluminescent bacterial growth assay

Upon infection of HeLa cells with *Stm*-lux, bacterial growth was followed over time by incubating the cells in the SpectraMax i3x plate reader at 37°C and measuring the bioluminescence every 15 minutes for 18 hours.

### Zebrafish husbandry

Zebrafish were handled in compliance with animal welfare regulations and maintained according to standard protocols (http://zfin.org). Fertilized embryos were maintained at 28°C and kept in egg water containing 60 μg/ml Instant Ocean Sea Salt (Sera, Heinsberg Germany). Zebrafish larvae were anesthetized with egg water containing 0.02% buffered 3-aminobenzoic acid ethyl ester (Tricaine, Sigma-Aldrich, Netherlands) for bacterial infection and imaging experiments.

### Zebrafish embryo toxicity test

Zebrafish embryos were manually dechorionated around 24 hours post fertilization (hpf) to exclude any protective effects from the egg shells. At 28 hpf, the compounds were added to the egg at concentrations ranging between 0.1-10 µM. At 120 hpf, the embryos were visually inspected using a Leica M205 FA stereo fluorescence microscope. The health of the embryos was scored with a maximum score of 5 (*i.e*., healthy), 1 point redrawn for no reaction to mechanical stimulation, oedema, tail curvature malformation, or cranial malformation, and a score of 0 for dead embryos without a heartbeat.

### Zebrafish embryo infection models

To test *Stm* compounds for efficacy *in vivo*, embryos that had already hatched from the egg at 48 hpf were collected and systemically infected with *Stm*-DsRed at 52 hpf by injecting 200 CFUs in the Duct of Cuvier as previously described [24]. Infected zebrafish embryos were given compounds at 54 hpf. The bacterial burden was quantified at 72 hpf by measuring fluorescence using the Leica M205 FA microscope. To test *Mtb* compounds for efficacy *in vivo*, dechorionated zebrafish embryos were systemically infected with *Mmar*-mWasabi at 28 hpf by injecting approximately 200 CFUs in the blood island (*i.e.*, caudal vein area), as previously described [24]. The compounds were added to the egg water at 30 hpf. The bacterial burden was determined at 144 hpf by fluorescence using the Leica M205 FA microscope and quantified using Fiji software [25].

### Statistical analyses

Statistical testing was performed using GraphPad Prism 9 (GraphPad Software, San Diego, CA, USA). Differences in the level of kinase inhibition between hit compounds and non-hit compounds were tested for statistically significant differences using Mann-Whitney tests. The results of intracellular infection and bacterial growth assays were tested for statistically significant differences between treatment and DMSO groups by performing Friedman tests for matched samples and Dunn’s multiple comparisons tests for *post-hoc* analysis. The results of zebrafish experiments were tested for statistically significant differences between treatment and control groups (either DMSO or untreated) by performing Kruskal-Wallis tests for independent samples and Dunn’s multiple comparisons tests for *post-hoc* analysis. Differences between groups resulting in *p*-values < 0.05 were considered statistically significant.

## Results

### Screening the PKIS library against intracellular *Stm* and *Mtb* identifies novel kinase inhibitors as candidates for host-directed therapy

In this study, a library of 827 PKIS kinase inhibitors was subjected to two consecutive screens to identify novel host-directed therapeutics with antimicrobial activity against intracellular *Stm* and *Mtb*. The results of the screen with *Stm*-DsRed-infected HeLa cells showed that two populations of infected cells could be discerned by flow cytometry, namely DsRed-dim and DsRed-bright (**Supplementary Figure 1a**). For *Mtb*, only one DsRed+ population was observed (**Supplementary Figure 1b**). As previously reported, the positive control H89, a PKA and Akt/PKB inhibitor known to reduce intracellular *Stm* bacterial burden, was found to diminish mainly the DsRed-bright population [9]. Since we observed the same phenomenon for several PKIS compounds, we set out to determine the contribution of the DsRed-dim and DsRed-bright populations of *Stm*-infected HeLa cells to the actual bacterial burden in order to choose the best outcome parameter. First, the presence of both populations was determined 1 h post infection and the DsRed-bright population was absent at this early timepoint (**Supplementary Figure 2a**). Next, after overnight incubation the DsRed-dim and DsRed-bright *Stm*-infected HeLa populations were sorted by FACS and lysed to determine the average CFUs/cell. DsRed-bright cells contained 142-times more viable bacteria than DsRed-dim cells (**Supplementary Figure 2b and c**). Combined, these results show that the DsRed-bright population comprises HeLa cells with replicating *Stm*, thus being most relevant as outcome parameter. The primary screen conducted with *Stm*-infected HeLa cells identified 82 inhibitors resulting in smaller DsRed-bright populations (z-score DsRed-bright < –2; **Supplementary Figure 1c and Supplementary Table 1**), of which 56 were not cytotoxic as determined by cellular counts by flow cytometry (z-score cell count > –3; **Supplementary Figure 1d**). The screen conducted with *Mtb*-infected MelJuSo cells identified 66 inhibitors (z-score DsRed+ < –2; **Supplementary Figure 1e**), of which 44 compounds did not result in cytotoxicity (z-score cell count > –3; **Supplementary Figure 1f**). From the 56 and 44 inhibitors of intracellular *Stm* and *Mtb*, respectively, 7 compounds were effective against both pathogens (**Supplementary Figure 1g**).

Rescreens were performed in order to select for compounds that consistently reduced the intracellular bacterial burden. For the rescreen a smaller set of PKIS compounds was used, comprising 201 compounds that were non-cytotoxic in the primary screen and found to be active, affecting the bacterial burden either positively or negatively (z-scores < –2 or > 2; **Supplementary Figures 1c and e**). For *Stm*, 14 PKIS hit compounds were identified in the rescreen, 11 of which also inhibited *Stm* in the primary screen and these were selected for further experiments (**Figure 1a-c**). For *Mtb*, 19 unique compounds were found to reduce the intracellular burden in the rescreen, 17 of which also inhibited *Mtb* in the primary screen. (**Figure 1d-f**). As one *Mtb* compound (*i.e*., GSK2289044B) could not be included in subsequent experiments because additional quantities were not available, 16 *Mtb*-effective compounds were selected for further experiments. In line with the cytotoxicity data from the primary screen, none of the *Stm* and *Mtb* hit compounds significantly affected the cell count (**Supplementary Table 1**). Ultimately, the screen resulted in 11 hit compounds against intracellular *Stm* and 16 hit compounds against intracellular *Mtb* that were further evaluated.

**Figure 1.**
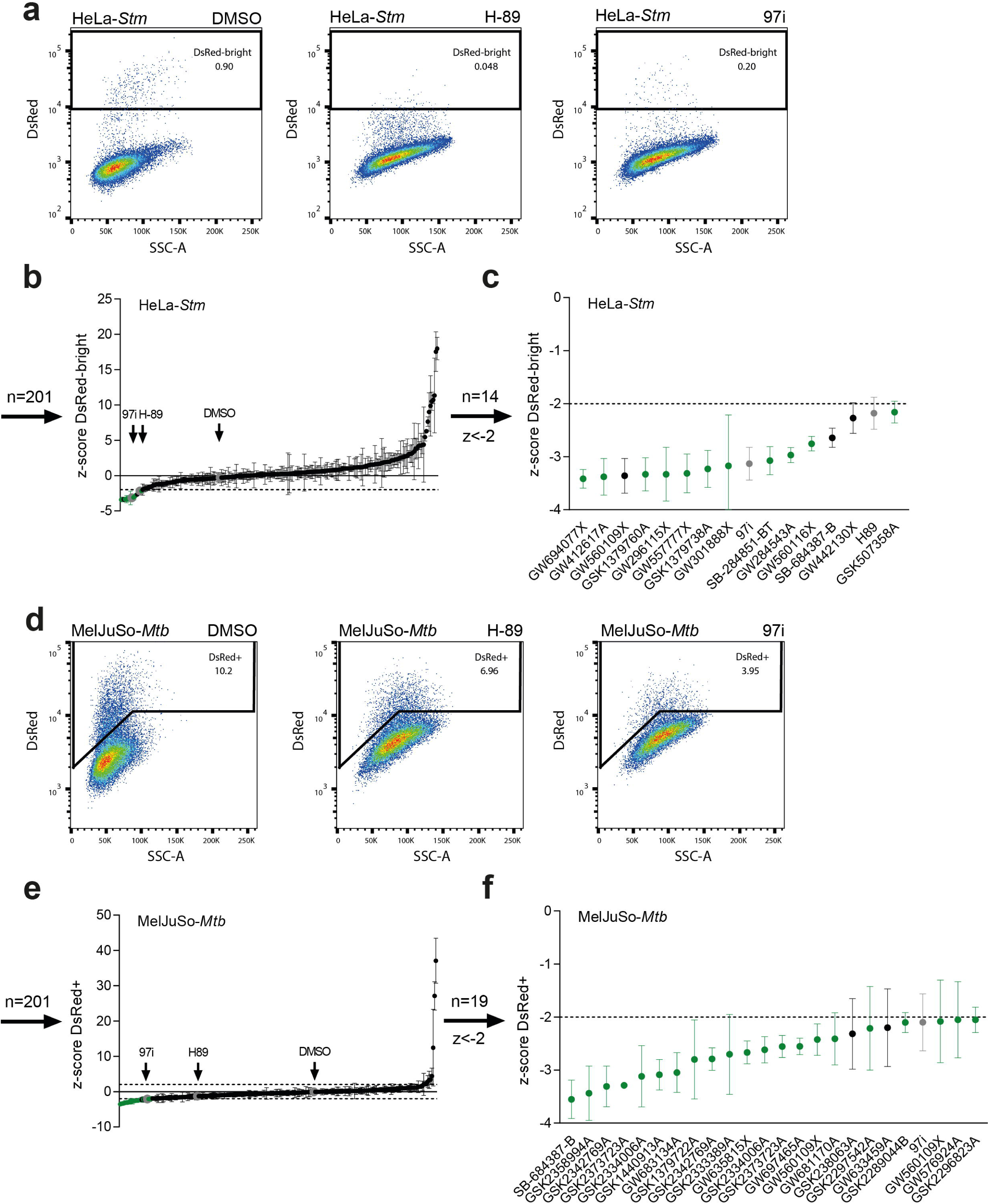
Identification of PKIS compounds inhibiting intracellular growth of *Stm* and *Mtb*. (**a**) Gating strategy for DsRed-bright *Stm*-infected HeLa cells to determine the percentage of infected cells. The negative control treated with DMSO and positive controls treated with H89 and 97i are depicted. (**b**) Rescreen of 201 PKIS compounds, 30 of which appear twice, to assess their impact on *Stm* bacterial burden, expressed as average z-scores of the DsRed-bright population. (**c**) Hit compounds with z-scores < –2. PKIS compounds that also reduced the bacterial burden in the primary screen are shown in green and others in black. (**d**) Gating strategy for DsRed+ *Mtb*-infected MelJuSo cells. (**e**) Re-screen on *Mtb*-infected MelJuSo cells. (**f**) Hit compounds with z-scores < –2 for *Mtb*. The screens were performed with three technical replicates, and error bars show standard deviations.

### Kinase profiling and structural analysis of *Stm* hit compounds identifies AAK1 as frequent target

Based on the published kinase inhibition profiles of the PKIS compounds, we determined which host kinases are likely important for host-pathogen interactions of *Stm*-infected cells [15,16]. The *Stm* hit compounds comprised 2 PKIS1 compounds and 9 PKIS2 compounds that have been profiled at 1 µM concentration against 203 and 392 wildtype kinases, respectively. For each kinase, compounds with > 50% kinase inhibition were counted. Commonly targeted kinases included MAP2K5 (6/9), PDGFRb (5/11), EphB6 (4/9) and AAK1 (5/9), which were also previously identified in a kinase gene silencing screen using *Stm*-infected HeLa cells [9] (**Figure 2a**). To ensure that the kinases were not simply common targets in the PKIS library (i.e., association bias), we compared the degree of inhibition of identified kinases between hit compounds and non-hit compounds (**Figure 2b**). The hit compounds exhibited greater inhibition in all identified targets, although this difference was only statistically significant for AAK1. Two pairs of *Stm* hit compounds were structurally related. The 2-anilino-4-pyrrolidinopyrimidines GSK1379760A and GSK1379738A only differed in their 2-anilino group, with GSK1379760A possessing two additional 3-, and 5-methoxyl groups), and while the compounds show limited overlap in their targets, both target JAK2 and AAK1 (**Figure 2c**). In addition, 4-anilino-quinolines GW557777X and GW560116X only differed in their 4-anilino group, with GW557777X possessing 2-methyl and 5-hydroxyl groups and GW560116X possessing 2-fluorine and 4-chlorine groups, and both target SRC, BLK, EphB6, KIT, PDGFRa, PDGFRb, RIPK2, ACTR2B, MAP2K5 and RSK4 (**Figure 2d**). Furthermore, compound GSK507358A is an inhibitor of all Akt isoforms and may therefore act in a similar way as H-89 [26]. Of note, two similar Akt-targeting compounds, GSK682037B and GSK562689A, also strongly reduced the *Stm*-bright population, but were excluded due to cytotoxicity (**Supplementary Table 1**).

**Figure 2.**
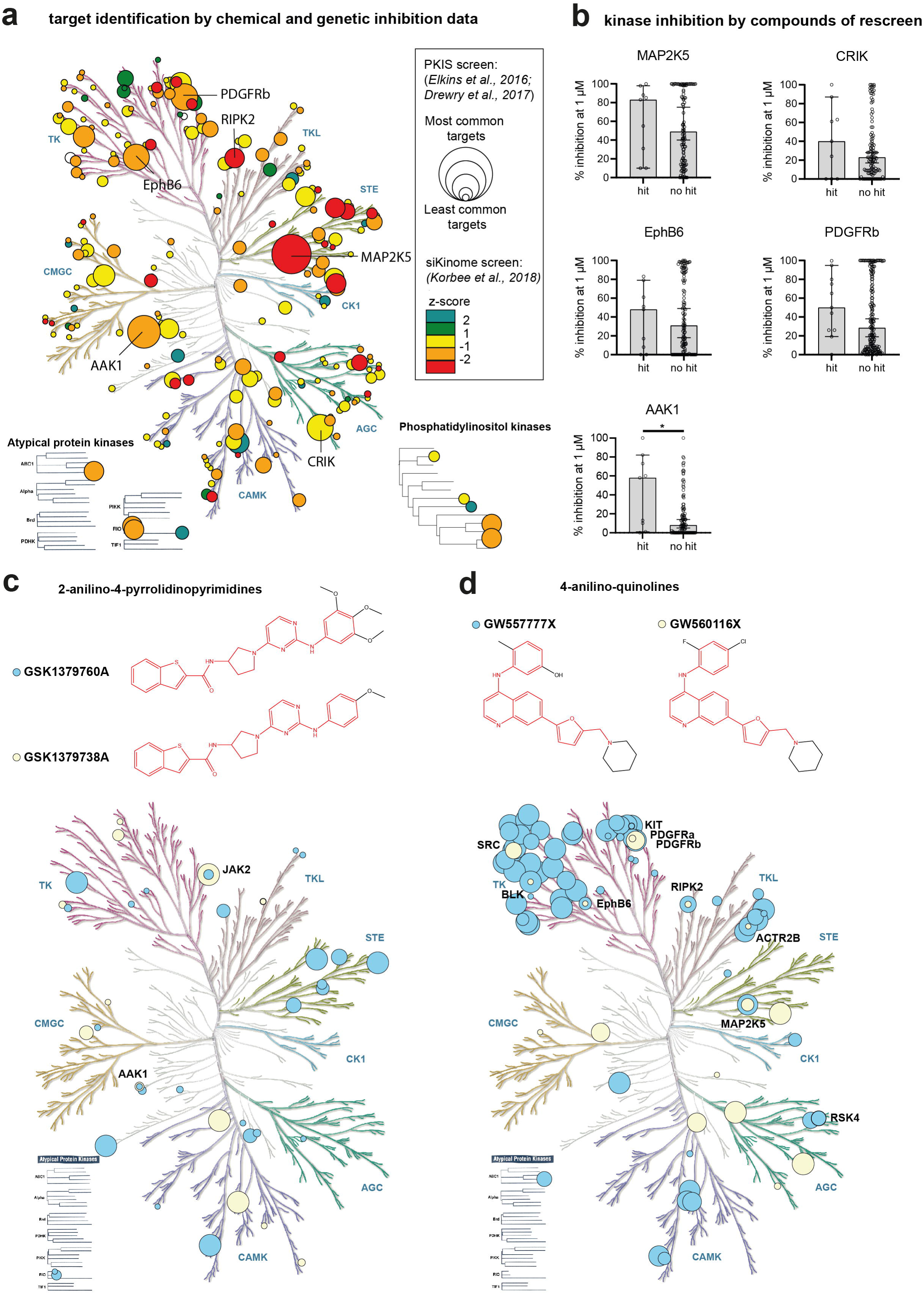
Kinase target profiling of *Stm* hit compounds. (**a**) The phylogenetic tree depicts host kinases that are relevant for HDT against intracellular *Stm* by combining data on chemical and genetic kinase inhibition. The size of each circle indicates the frequency at which a kinase was targeted by *Stm* hit compounds, while the color of the circle represents the effect of genetic inhibition on *Stm* burden in the HeLa-*Stm* model, as determined previously by our research group [9]. (**b**) Comparison of kinase inhibition between PKIS compounds that reduced intracellular *Stm* (hit) and compounds that did not (no hit). Statistically significant differences are indicated by \**p* < 0.05. (**c**) Target comparison between two structurally related 2-anilino-4-pyrrolidinopyrimidine hit compounds. (**d**) Target comparison between two structurally related 4-anilino-quinoline hit compounds.

### Morpholino-imidazo/triazolo-pyrimidinones targeting phosphatidylinositol 3-kinases and 2-aminobenzimidazoles targeting receptor tyrosine kinases are effective against intracellular *Mtb*

The *Mtb* hit compounds comprised 1 PKIS1 compound, 15 PKIS2 compounds and 1 compound that is part of both sets. Commonly targeted kinases were ABL1 (7/17), EphA1 (6/15), EphB6 (6/15), RET (9/17), MAP3K19 (7/15), MAP2K5 (6/15), CRIK (6/15), PIK3CB (8/15), PIK3CD (7/17) and VPS34 (8/15) (**Figure 3a**). Genetic inhibition of ABL1, PIK3CB, PIK3CD and VPS34 was previously found to reduce bacterial burden in the MelJuSo-*Mtb* intracellular infection model [9]. Again, we compared the degree of inhibition of the identified kinase targets between the hit compounds and other compounds of the PKIS library and found that RET, PIK3CB, PIK3CD and VPS34 were inhibited significantly more by hit compounds (**Figure 3b**). Among compounds that inhibited phosphatidylinositol 3-kinases PIK3CB, PIK3CD and VPS34, there was a large group of structurally related morpholino-imidazo/triazolo-pyrimidinones (**Figure 3c**). These compounds were specific for PIK3CB, PIK3CD and VPS34 and did not target other phosphatidylinositol kinases (**Figure 3d**). In addition, a group of 2-aminobenzimidazoles was found to target receptor tyrosine kinases ABL1, RET, TRKA, and to a lesser extent BLK. RET is inhibited by all 2-aminobenzimidazoles included in the PKIS library, regardless of their effect on *Mtb* bacterial burden, and might therefore be a false-positive target (**Supplementary Table 1**) [16].

**Figure 3.**
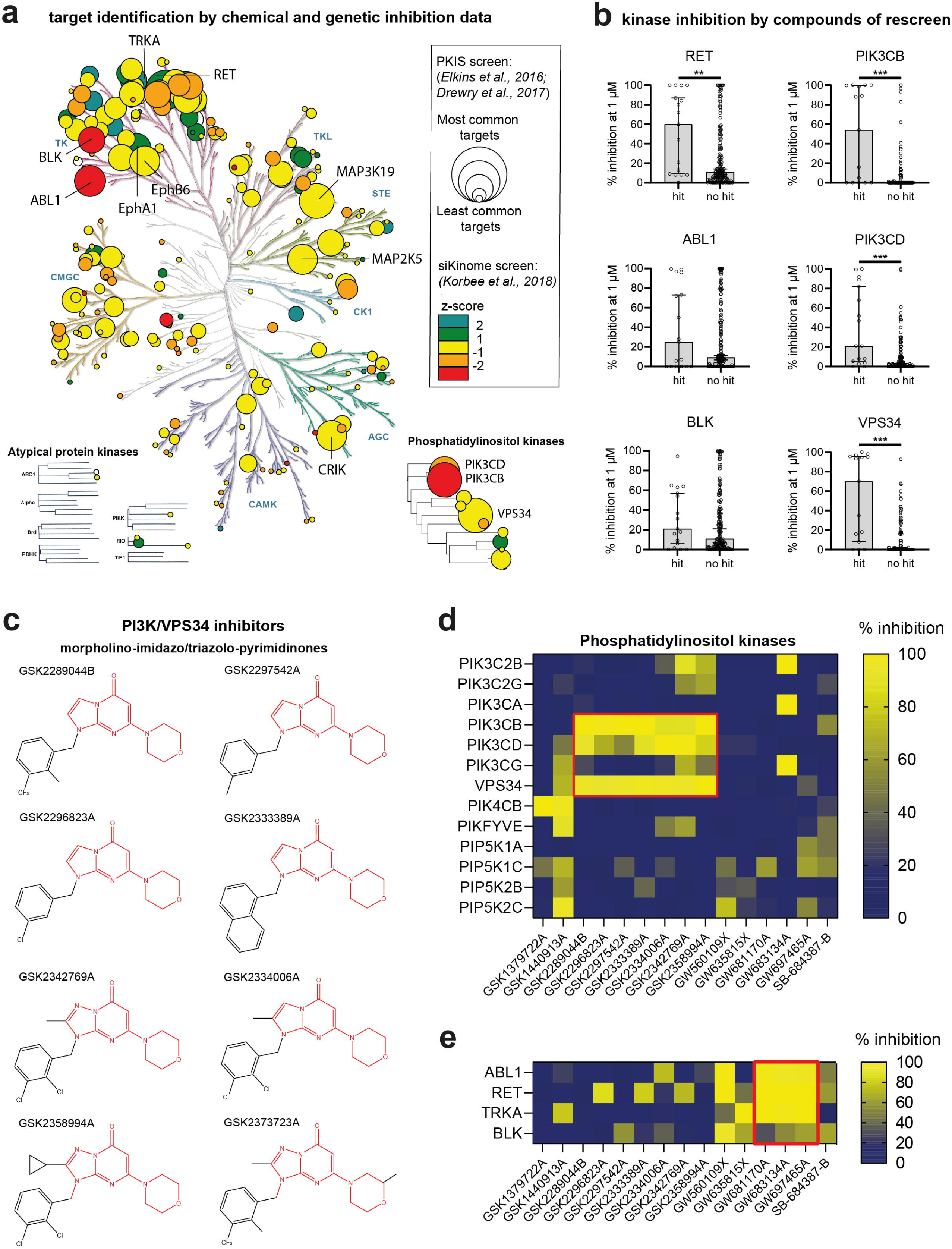
Kinase target profile of *Mtb* hit compounds. (**a**) The phylogenetic tree depicts host kinases that are relevant for HDT against intracellular *Mtb* by combining data on chemical and genetic kinase inhibition. The size of each circle indicates the frequency at which a kinase was targeted by *Mtb* hit compounds, while the color of the circle represents the effect of genetic inhibition on *Mtb* burden in the MelJuSo-*Mtb* model, as determined previously by our research group [9] (**b**) Comparison of kinase inhibition between PKIS compounds that reduced intracellular *Mtb* (hit) and compounds that did not (no hit). Statistically significant differences are indicated by \*\**p* < 0.01 and \*\*\**p* < 0.001. (**c**) Chemical structures of structurally-related morpholino-imidazo/triazolo-pyrimidinones with shared phosphatidylinositol 3-kinase targets. (**d**) Inhibition of phosphatidyl inositol kinases by PKIS2 compounds [16]. Red square indicates targets of morpholino-imidazo/triazolo-pyrimidinones. (**e**) Inhibition of selected protein kinases that were previously identified as HDT targets for *Mtb* [9,32]. Red square indicates targets of 2-aminobenzimidazoles.

### All *Stm* hit compounds and the majority of *Mtb* hit compounds reduce bacterial counts, acting through a host-directed manner

Validation of the efficacy of hit compounds in CFU assays showed that all *Stm* hit compounds effectively reduced the intracellular bacterial burden of *Stm*-infected HeLa cells, with 8 out of 11 compounds that showed > 90% inhibition (**Figure 4a**). The *Mtb* hit compounds showed a more modest reduction in intracellular bacterial load with 11 compounds reducing *Mtb* bacterial load > 25% and 2 compounds (*i.e.*, GW560109X and GW635815X) leading to a reduction greater than the 97i positive control (**Figure 4b**). The safety of the compounds was validated further by an LDH-release cytotoxicity assay. *Stm* compound GW301888X (**Figure 4c**) and *Mtb* compound GW560109X (**Figure 4d**) reduced HeLa and MelJuSo cell viability by 15% and 17%, respectively, while no noteworthy loss in viability was observed for the other compounds. Finally, we assessed whether the selected *Stm* and *Mtb* hit compounds exerted any direct antibacterial effects on planktonic bacteria to confirm that the compounds act as HDTs. For *Stm*, none of the compounds showed direct effects in planktonic *Stm* culture (**Figure 4e**). For *Mtb*, compounds GW560109X and SB-684387-B showed some direct effect and reduced the bacterial concentration in planktonic cultures by 27% and 18%, respectively, after 13 days of incubation (**Figure 4f**).

**Figure 4.**
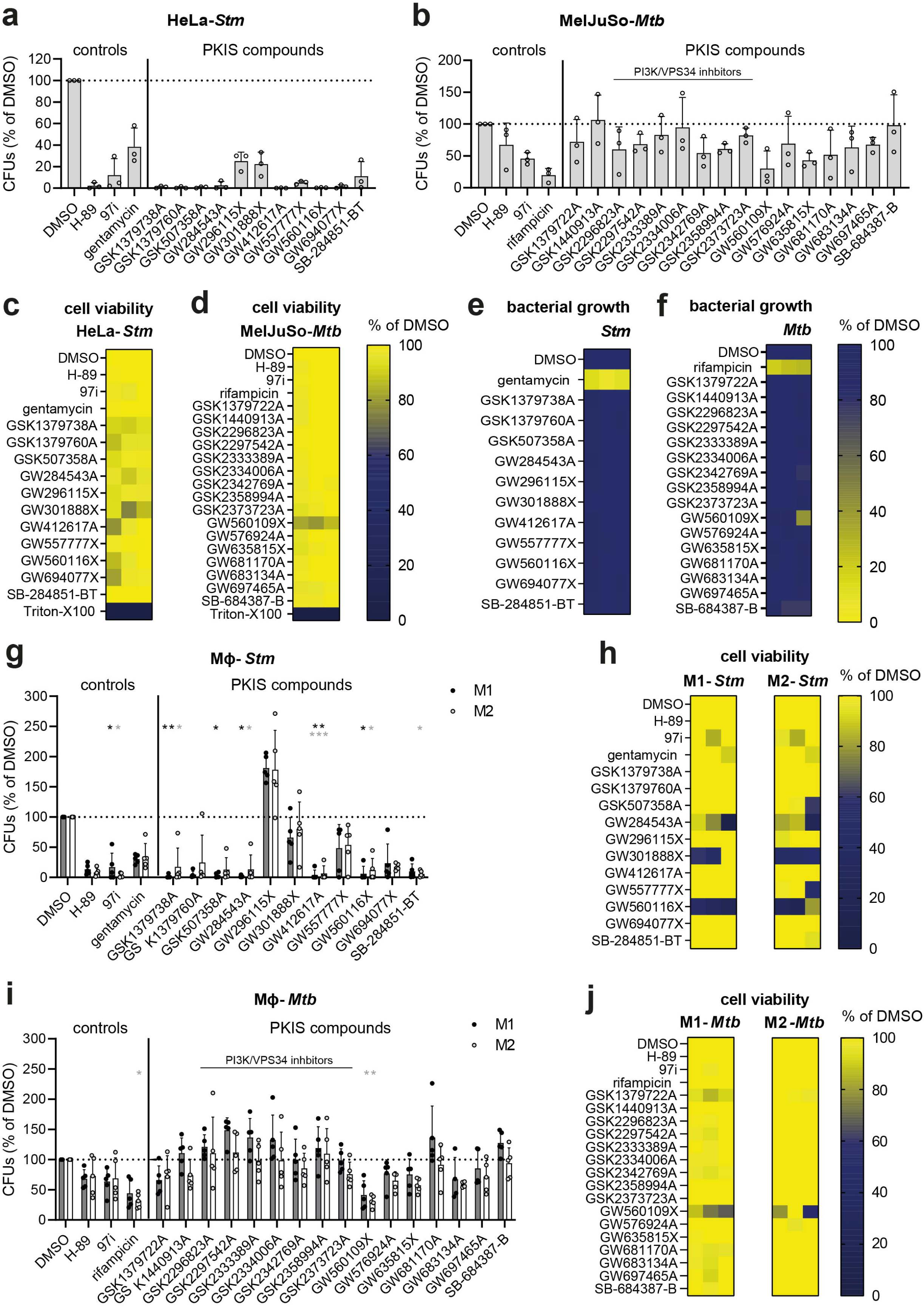
Validation of PKIS hit compounds in cell lines and primary human macrophages. (**a-b**) The efficacy of the hit compounds was validated in CFU assays using lysates from *Stm*-infected HeLa cells (**a**) and *Mtb*-infected MelJuSo cells (**b**). The bacterial burden is expressed as a percentage of CFUs compared to the DMSO control. (**c-d**) Compound safety was assessed using an LDH-release assay using supernatant from *Stm*-infected HeLa cells (**c**) and *Mtb*-infected MelJuSo cells (**d**), with cell viability expressed as a percentage of the DMSO control and with 1% Triton-X100-treated cells corresponding to 0%. (**e-f**) To assess whether hit compounds act as antibiotics or host-directed therapeutics, direct antimicrobial effects were evaluated in cell-free cultures of *Stm* (**e**) and *Mtb* (**f**). The turbidity of the bacterial suspensions, as measured by absorbance at OD_600_, is given as a percentage of the DMSO control. (**g**) The efficacy of the hit compounds was validated in CFU assays using lysates from *Stm*-infected M1 (black circles, grey bars) and M2 (white circles, open bars) primary human macrophages. (**h**) An LDH-release assay was performed using supernatant from *Stm*-infected macrophages. (**i**) The efficacy of the hit compounds was validated in CFU assays using lysates from *Mtb*-infected M1 (black circles) and M2 (white circles) primary human macrophages. (**h**) An LDH-release assay was performed using supernatant from *Mtb*-infected macrophages.

### Many identified hit compounds are also effective against *Stm* or *Mtb* infected primary human macrophages, while phosphatidylinositol 3-kinase inhibitors are only effective against *Mtb*-infected MelJuSo cells

Macrophages play a critical role during *Stm* and *Mtb* infections, both as part of the immune response and as the preferred host cell type for the bacteria. Therefore, hit compounds were tested in *Stm*-infected and *Mtb*-infected primary human macrophages. CD14+ monocytes were isolated from PBMCs of healthy human donors and the cells were cultured in the presence of GM-CSF and M-CSF to generate M1 and M2 macrophages, respectively (**Supplementary Figure 4a-c**). The two types of macrophages were clearly distinguishable by expression of CD11b, CD163 and CD14, in accordance with previous publications (**Supplementary Figures 4d-e**) [19]. The majority of the compounds that were effective in *Stm*-infected HeLa cells were also highly effective in both types of *Stm*-infected macrophages (**Figures 4g**). However, compounds GW296115X, GW301888X and GW557777X showed reduced effectiveness in *Stm*-infected macrophages. In contrast to HeLa cells, in which none of the compounds were found to be cytotoxic, compounds GW284543A (M1: 58.4%; M2: 64.3%), GW301888X (M1: 68.8%; M2: 47.9%) and GW560116X (M1: 34.4%; M2: 41.3%) reduced cell viability of primary human macrophages (**Figure 4h**). Testing the *Mtb* hit compounds on *Mtb*-infected macrophages showed that many compounds were as effective as in *Mtb*-infected MelJuSo cells, with compound GW560109X being most effective in both cell types (**Figures 4i**). In addition, the moderate cytotoxicity of compound GW560109X observed in MelJuSo cells was also observed in primary human macrophages (**Figure 4j**). In contrast, the group of morpholino-imidazo/triazolo-pyrimidinones acting on PI3K/VPS34 and compound GW681170A were effective in *Mtb*-infected MelJuSo cells, but not in primary human macrophages (**Figures 4b and 4i**).

### *Stm* hit compounds act readily by inhibiting intracellular bacterial growth and several display a high therapeutic index *in vitro*

After having validated the effectiveness of many *Stm* hit compounds in infected primary human macrophages, we performed dose-titration experiments on *Stm*-infected HeLa cells for the seven most efficacious compounds to determine their therapeutic index, which is important for their potential for further progression. In order to follow bacterial growth over time for 18 hours, we performed dose-titration experiments using a bioluminescent *Stm*-lux strain (**Figure 5a**). After a short lag phase during which the bacteria were controlled by the host cells, *Stm* started to grow exponentially. At their effective concentration, all compounds acted readily by inhibiting the bacterial growth rate. The minimal inhibitory concentration (MIC)-values of the compounds ranged from 1.8-14 µM (**Table 1**). Dose-response curves were created to determine the *in vitro* potency and safety of the compounds (**Figure 5b**). The IC_50_-values ranged from 1.0-6.5 µM and selectivity indexes from 6.5 – 107 (**Table 1**). Some compounds showed some cytotoxicity at the tested concentrations, with GW284543A being the most cytotoxic compound with an LD_50_-value of 24 µM. Interestingly, compound GW560116X showed no cytotoxicity in HeLa cells, in contrast to primary human macrophages for which significant cytotoxicity was observed at 10 µM concentration (**Figure 4h**).

**Figure 5.**
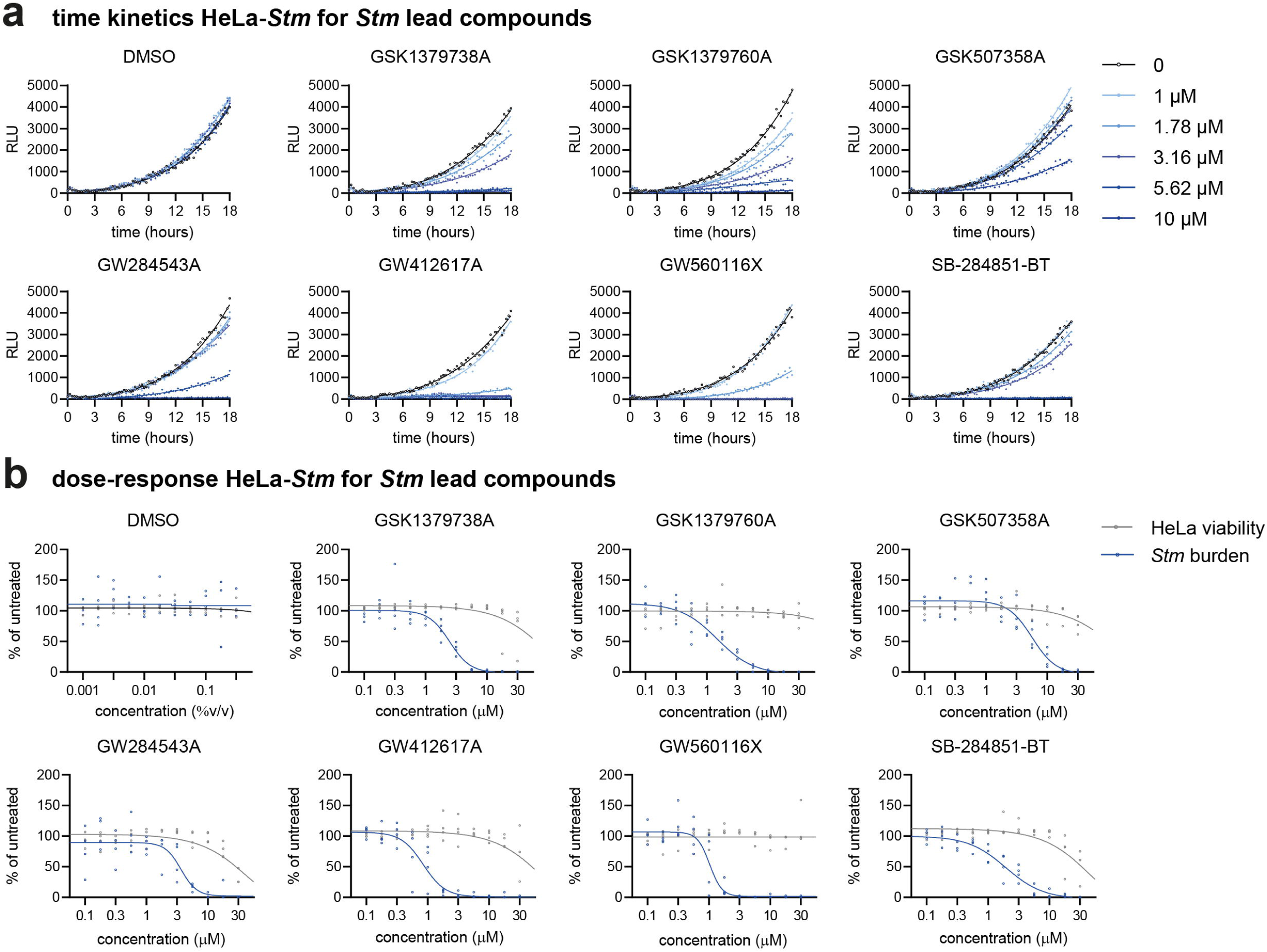
Time kinetics and dose-response relationship *Stm* lead compounds in *Stm*-infected HeLa cells. (**a**) HeLa cells were infected with bioluminescent *Stm*-lux, treated with *Stm* lead compounds or DMSO at different concentrations and followed over time by measuring emitted light, expressed as relative light units (RLU). For DMSO, equal %v/v was used for treatment. Gompertz growth curves were fitted by non-linear regression. (**b**) Bioluminescence, shown in blue, was measured for HeLa cells infected with *Stm*-lux after 18 h of treatment with *Stm* lead compounds at indicated concentrations. RLUs were normalized to the untreated control. Four parameter logistic regression was performed to show the dose-response relationship between compounds and inhibition of *Stm* growth and to determine the half maximal inhibitory concentration (IC_50_) of compounds. In addition, the cell supernatant was used to determine host cell viability, shown in grey, using LDH-release assays. Again, four parameter logistic regression was performed to the determine the half maximal lethal dose (LD_50_) of compounds. The data comprises 4 independent technical replicates.

**Table 1.** *In vitro* efficacy (E_max_), minimal inhibitory concentration (MIC), potency (IC_50_), safety (LD_50_) and therapeutic index (TI; ratio between IC_50_ and LD_50_) of *Stm* lead compounds against intracellular *Stm*. Numbers between brackets show the 95% confidence interval. Some values could not be determined (ND).

| Compound | E<sub>max</sub> (% inh.) | MIC (µM) | IC<sub>50</sub> (µM) | LD<sub>50</sub> (µM) | TI |
| --- | --- | --- | --- | --- | --- |
| GSK1379738A | 100 (92-100) | 6.7 (3.2-10) | 2,5 (2.0-3.1) | 45 (34-116) | 18 |
| GSK1379760A | 100 (93-100) | 6.1 (1.6-11) | 1,8 (1.4-2.3) | 161 (ND) | 107 |
| GSK507358A | 100 (87-100) | 14 (6.7-21) | 6.5 (4.9-9.1) | 49 (ND) | 8.6 |
| GW284543A | 98 (85-100) | 7.8 (3.8-12) | 3,2 (2.2-5.9) | 24 (21-27) | 6.5 |
| GW412617A | 99 (91-100) | 1.8 (0.1-3.5) | 1.1 (0.7-1.2) | 33 (28-50) | 37 |
| GW560116X | 99 (90-100) | 1.9 (0.5-3.4) | 1,0 (0.9-1.3) | ND | ND |
| SB-284851-BT | 100 (93-100) | 6.7 (3.2-10) | 2.0 (1.5-2.7) | 26 (22-32) | 14 |

### Structurally related *Stm* hit compounds GSK1379738A and GSK1379760A exhibited significant effectiveness in infected zebrafish embryos

Finally, we determined whether the seven most effective *Stm* compounds and six most effective *Mtb* compounds were safe and efficacious *in vivo* using zebrafish embryo models. In vivo toxicity was determined by administrating the compounds directly to the water of dechorionated zebrafish embryos (**Figure 6a**). After 4 days the health of the embryos was scored based on their mortality rate and drug-induced side effects (**Figure 6b**). For the *Stm* compounds, only GSK1379738A resulted in some toxicity, causing oedema (4/20) at 10 µM concentration (**Figure 6c**). *Mtb* compound GW683134A caused oedema and cranial malformations in the majority of embryos starting at 0.3 µM and 1 µM, respectively (**Figure 6d**). In addition, GW697465A caused oedema in a large proportion of the embryos starting at 1 µM. The other *Mtb* compounds were considered non-toxic *in vivo*. Overall, mortality was low even for the toxic compounds at the tested concentrations.

**Figure 6.**
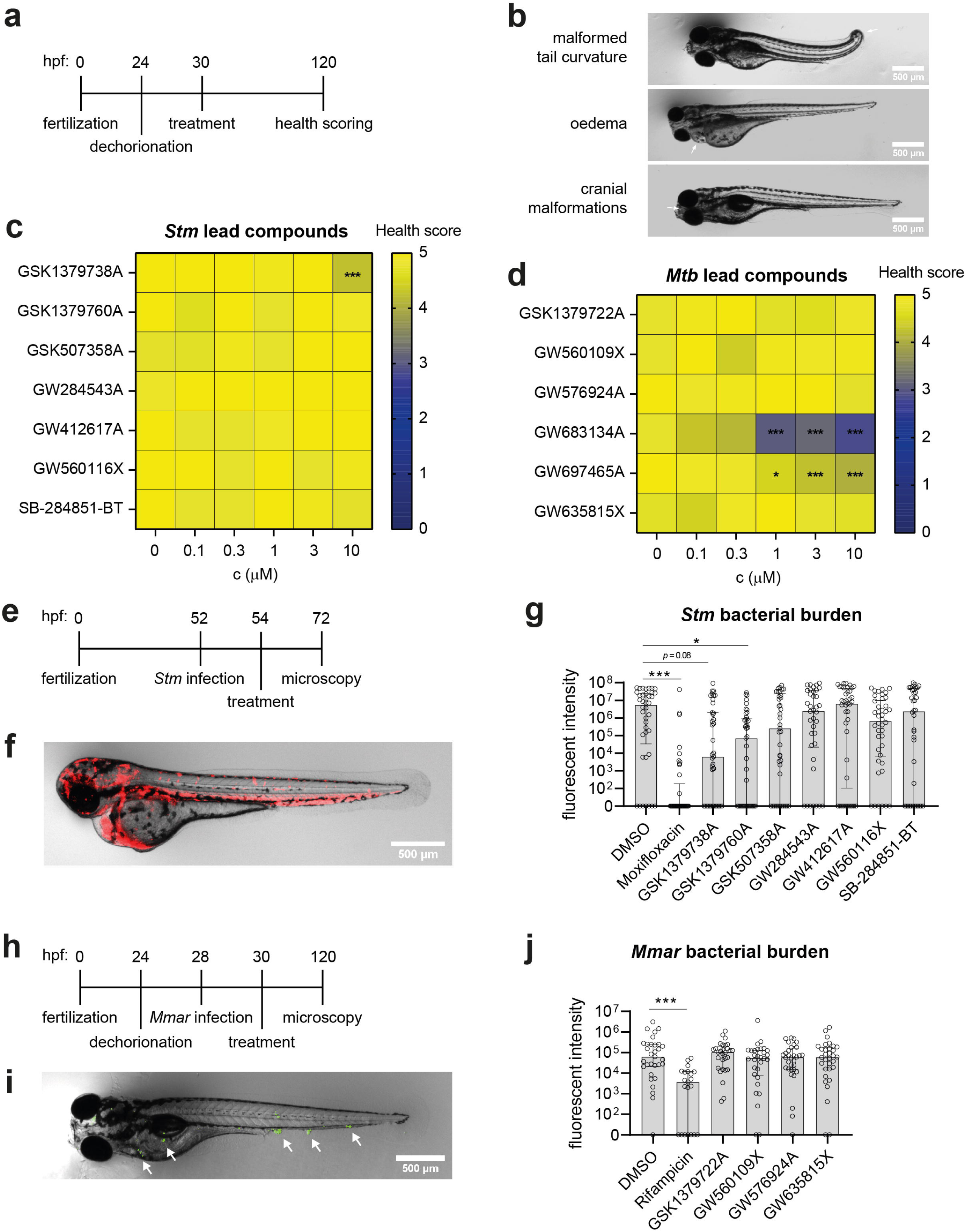
Testing of *Stm* and *Mtb* lead compounds *in vivo* in zebrafish embryo models. (**a**) Schematic representation of the zebrafish embryo toxicity model. Embryos were visibly inspected and scored for the health with a score of 5 representing healthy embryos, 0 representing dead embryos, and scores in between representing embryos with one to four of the following conditions: malformed tail curvature, oedema, cranial malformations, no response to physical stimulation. (**b**) Representative images of embryos with malformations. (**c-d**) Zebrafish embryos were treated with *Stm* (**c**) and *Mtb* (**d**) lead compounds at different concentrations. Each square of the heat map represents the average health score of 20 embryos. (**e**) Schematic representation of the zebrafish embryo *Stm*-infection model. Embryos were infected with ∼200 CFUs 52 hours post fertilization (hpf). (**f**) Representative image of *Stm*-infected zebrafish embryos treated with DMSO 72 hpf. (**g**) *Stm*-infected zebrafish were treated with 0.1%v/v DMSO, 1 µM moxifloxacin, 3 µM GSK1379738A and other compounds at 10 µM. Each group comprises 38-42 embryos. (**h**) Schematic representation of the zebrafish embryo *Mmar*-infection model. Embryos were infected with ∼200 CFUs 28 hpf. (**i**) Representative image of *Mmar*-infected zebrafish embryos treated with DMSO 120 hpf. (**j**) *Mmar*-infected embryos were treated with 0.1% DMSO, 200 µM rifampicin and other compounds at 10 µM. Each group comprises 21-32 embryos. Statistically significant differences are indicated by \**p* < 0.05 and \*\*\**p* < 0.001.

The efficacy of the compounds was tested in zebrafish embryo infection models. Zebrafish embryos were injected with 200 CFUs *Stm*, treated with the *Stm* compounds, and the bacterial burden was quantified 20 hours later (**Figures 6e-f**). Of note, compound GSK1379738A was used at 3 µM instead of 10 µM, as used for the other compounds, to prevent toxicity. Compounds GSK1379738A, GSK1379760A and GSK507358A were highly effective, resulting in 870, 77 and 21-times lower median bacterial burdens, respectively, as compared to DMSO-treated controls (**Figure 6g**). Due to variation between individuals, only GSK1379760A led to a statistically significant reduction. Next, zebrafish embryos were injected with approximately 200 CFUs *Mmar*, an established model of human *Mtb* infections [27,28], to test the *in vivo* effectiveness of the non-toxic *Mtb* compounds GSK1379722A, GW560109X, GW576924A and GW635815X upon 4 days of treatment (**Figure 6h**). Granuloma formation was clearly observed in infected embryos (**Figure 6i**). Rifampicin was used as a positive control and resulted in a 29-times lower median bacterial burden compared to the DMSO control (**Figure 6j**). However, none of the *Mtb* compounds were able to reduce the *Mmar* bacterial burden significantly. Taken together, most PKIS compounds with activity against intracellular *Stm* and *Mtb in vitro* were found safe *in vivo*, and two structurally related 2-anilino-4-pyrrolidinopyrimidines, GSK1379738A and GSK1379760A, were highly effective at reducing the *Stm* bacterial burden in vivo, showing the potential of this chemotype to use as HDT during *Stm* infections.

## Discussion

In response to the ongoing rise in antimicrobial resistance and limited effectiveness of antibiotics against intracellular bacterial species, we aimed to identify new HDTs that may be used as adjunctive or alternative treatments to classical antibiotics. Starting with 827 ATP-competitive kinase inhibitors from the PKIS library, we performed two consecutive screens in intracellular HeLa-*Stm* and MelJuSo-*Mtb* infection models. This resulted in 11 hit compounds that stimulate killing of intracellular *Stm* and 16 hit compounds that target intracellular *Mtb*, all of which were non-cytotoxic and acted in a host-directed manner. There was no overlap between *Stm* and *Mtb* hit compounds or their kinase targets, suggesting that these intracellular pathogens likely depend on very different signaling pathways to persist intracellularly. Importantly, all *Stm* and multiple *Mtb* hit compounds were also effective in primary human macrophages, which play a key role in the pathogenesis of these bacteria, both as host cells and effector immune cells [29,30]. Moreover, we were able to demonstrate that two related *Stm* compounds, GSK1379738A and GSK1379760A, effectively reduced the *Stm* bacterial burden in infected zebrafish embryos.

When studying kinase inhibition profiles of the PKIS hit compounds, those directed against *Mtb* were clearly dominated by a group of morpholino-imidazo/triazolo-pyrimidinones targeting PIK3CB, PIK3CD and VPS34. From these, PIK3CB and PIK3CD are most likely to be important for intracellular *Mtb* persistence in MelJuSo cells based on previously published genetic kinase inhibition data [9], in which knockdown of PIK3CB and PIK3CD, but not VPS34 reduced the intracellular bacterial burden of *Mtb*-infected MelJuSo cells. The higher efficacy of the morpholino-imidazo/triazolo-pyrimidinones in MelJuSo cells as compared to primary human macrophages may be explained by presence of a gain-of-function mutation of the NRAS gene at c.182A>T in MelJuSo cells, resulting in constitutively increased activity of PI3K [31]. PIK3CB and PIK3CD phosphorylate phosphatidylinositol 4,5-bisphosphate to create phosphatidylinositol 3,4,5-trisphosphate at the plasma membrane, which provides a binding site for many cytosolic proteins including Akt. Previous studies have shown that the PI3K/Akt pathway is important for intracellular persistence of both *Stm* and *Mtb* [6–8]. In this study, PI3K activity clearly facilitated persistence of *Mtb* in infected MelJuSo cells, but seemingly had no effect in *Mtb*-infected primary human macrophages or *Stm*-infected HeLa cells. Vice versa, inhibition of Akt by H-89 and GSK507358A had limited effect on *Mtb*-infected MelJuSo cells, but potently inhibited *Stm* in infected HeLa cells or primary human macrophages. Thus our results show that while PI3K/Akt kinases are important in both *Stm* and *Mtb* infection, the exact kinases involved in bacterial persistence are specific to both pathogen and host cell.

In addition to compounds that target the PI3K/Akt pathway, we identified *Mtb* hit compounds targeting receptor kinase receptors BLK, ABL1 and TRKA (also known as NTRK1), all of which have previously been identified as HDT targets against *Mtb* [6, 9–13,32]. Best known in this regards is ABL1 as the main target of imatinib, which is currently tested in clinical trials as adjunctive therapy together with an antibiotic regiment of rifabutin and isoniazid against drug-sensitive TB [14]. Compound GW560109X, although structurally unrelated to imatinib, is a strong ABL1 inhibitor and was most effective against intracellular *Mtb* from all PKIS compounds tested, both in infected MelJuSo cells and primary human macrophages [16]. The *Stm* hit compounds were part of diverse chemotypes with only a few targets in common, including AAK1, an enzyme involved in clathrin-mediated endocytosis. Inhibitors of AAK1 have been pursued as broad-spectrum antivirals that block viral entry. In addition, a recent paper has demonstrated that AAK1 mediates internalization of LPS, and blocking AAK1 prevents lethality during bacterial sepsis in mice [33].

Seven *Stm* lead compounds and six *Mtb* lead compounds were tested *in vivo* in zebrafish embryo models, of which Stm compounds GSK1379738A and GSK1379760A were highly effective. These compounds are structurally closely related, showing that this chemotype is interesting to further optimize for use as HDT. Possibly, GSK1379738A and GSK1379760A act by targeting JAK2 or AAK1, which are shared kinase targets. In addition, the Akt inhibitor GSK507358A also reduced the *Stm* bacterial burden by 20-fold, but this did not reach statistical significance due to the interindividual variation.

While the zebrafish model was successfully employed to confirm *Stm* compounds, none of the *Mtb* compounds showed effectiveness in the zebrafish TB model. One explanation could be that a closely related but different pathogen, Mmar, is used in this model. In addition, there may have been limitations in the way the *in vivo* experiments were performed. All compounds were given to zebrafish embryos by immersion in compound-supplemented egg water, which is the most commonly used method. However, recent studies have shown that some drugs may show limited absorption by the embryos depending on their lipophilicity [34,35]. This may also explain why the *in vitro* cytotoxicity data based on LDH-release assay was not found predictive for in vivo toxicity. For example, 10 µM *Stm* lead compound GW560116X and 10 µM *Mtb* lead compound GW560109X caused cell death when administered to primary human macrophages *in vitro*, but did not result in any side effects in zebrafish embryos. In future experiments, the route of administration of HDTs in may have to be optimized depending on their lipophilicity.

In summary, we identified morpholino-imidazo/triazolo-pyrimidinones targeting PIK3CB and PIK3CD as interesting compounds against intracellular *Mtb* and found that their effect was cell type-specific, showing reduced activity in primary human macrophages as compared to the MelJuSo cell line. Moreover, we identified 2-aminobenzimidazoles targeting receptor tyrosine kinases BLK, ABL1 and TRKA as effective against intracellular *Mtb*. Interestingly, these kinases were previously found as targets for HDT against *Mtb* [9]. For *Stm*, we identified 7 compounds that were highly potent in both *Stm*-infected HeLa cells and primary human macrophages, resulting in near clearance of the intracellular bacterial burden. Compound GSK1379760A, a 2-anilino-4-pyrrolidinopyrimidine, was found to be the most potent kinase inhibitor against intracellular *Stm*, showing a therapeutic index of >107-fold *in vitro* and reducing the bacterial burden by 77-fold in zebrafish embryos in vivo. Another 2-anilino-4-pyrrolidinopyrimidine, compound GSK1379738A, was effective against *Stm*, both *in vitro* and *in vivo*, which demonstrates the potential of this chemotype for use as HDT. This study furthermore demonstrates the capacity of the exploited pipelines to identify potent HDTs that are translatable and well tolerated *in vivo*. Further studies are required to demonstrate the potential of GSK1379760A as adjunctive or alternative treatment to antibiotics, including its ability to treat bacterial infection *in vivo* in mammals and humans.

## Acknowledgements

We thank Beatriz Urones Ruano, Pablo Castañeda Casado, Isabel Camino and Joel Lelievre from GlaxoSmithKline for their extensive collaboration and providing insightful information about the PKIS compounds. Their support enabled us to carry out comprehensive and in-depth analysis of these compounds. Furthermore, we thank prof. William Zuercher, prof. David Harold Drewry and colleagues from University of Chapel Hill, North Carolina for providing us with additional quantities of selected compounds from the Published Kinase Inhibitor Set which enabled us to conduct our experiments. Finally, we thank Lennert Janssen, Virginie Stevenin and prof. Jacques Neefjes from Cell & Chemical Biology department of the Leiden University Medical Center for providing the *Stm*-lux strain.

## Supplementary data

Supplementary Table 1 Raw data and z-scores obtained from flow cytometry screens of PKIS compounds.

Supplementary Figure 1 Primary screen of PKIS compounds affecting *Stm* and *Mtb* intracellular burden. (**a**) Gating strategy for DsRed+ and DsRed-bright *Stm*-infected HeLa cells. H89-treated cells illustrate that the DsRed-bright population may be reduced without reducing the DsRed+population. (**b**) Gating strategy for DsRed+ *Mtb*-infected MelJuSo cells. (**c**) Primary screen of 827 PKIS compounds to assess their impact on *Stm* bacterial burden, expressed both as average z-scores of the DsRed-bright population. Compounds with z-scores < –2 or > 2 are shown in green and red, respectively. (**d**) Cytotoxicity of compounds was determined by determining z-scores of HeLa cell counts. Compounds with a z-score > –3 were considered non-cytotoxic and included. (**e**) Similar to (**c**), with z-scores representing intracellular *Mtb* burden. (**f**) Similar to (**d**), with z-scores representing MelJuSo cell counts. The screens comprise three replicates for the PKIS compounds and error bars show the standard deviations. (**g**) The VENN diagram shows the overlap between the hit compounds from the HeLa-*Stm* and MelJuSo-*Mtb* screens.

Supplementary Figure 2 Comparison DsRed-dim and DsRed-bright *Stm*-infected HeLa cells. (**a**) HeLa cells were fixed either 1 h or 24 h after infection with *Stm*-DsRed to determine differences in the presence of both DsRed populations. (**b**) HeLa cells were selected for FACS sorting based on gates for single cells, size and DsRed fluorescence. (**c**) The DsRed-dim and DsRed-bright populations were sorted and lysed to determine the intracellular bacterial burden by CFU count.

Supplementary Figure 3 Generation, morphology and phenotype of primary human macrophages. (**a**) Representative histograms showing CD14 expression in PBMCs and CD14-MACS sorted monocytes. (**b**) Enrichment of CD14^+^ cells before and after MACS sorting. (**c**) Morphology of M1 and M2 macrophages. (**d**) Representative histograms showing distinct expression patterns of CD11b, CD163 and CD14 by M1 and M2 macrophages. (**e**) Expression of CD11b, CD163 and CD14 by M1 and M2 macrophages was tested for differences using Wilcoxon matched-paired signed rank tests. Statistically significant differences are shown by **p < 0.01.

## References

1. Stanaway JD, Parisi A, Sarkar K, Blacker BF, Reiner RC, Hay SI, et al. The global burden of non-typhoidal salmonella invasive disease: a systematic analysis for the Global Burden of Disease Study 2017. Lancet Infect Dis (2019) 19(12):1312–1324. doi:10.1016/S1473-3099(19)30418-9

2. James SL, Abate D, Abate KH, Abay SM, Abbafati C, Abbasi N, et al. Global, regional, and national incidence, prevalence, and years lived with disability for 354 diseases and injuries for 195 countries and territories, 1990-2017: a systematic analysis for the Global Burden of Disease Study 2017. The Lancet (2018) 392(10159):1789–1858. doi:10.1016/S0140-6736(18)32279-7

3. Houben RMGJ, Dodd PJ. The Global Burden of Latent Tuberculosis Infection: A Re-estimation Using Mathematical Modelling. PLoS Med (2016) 13(10):e1002152.

4. Organization WH. Global Tuberculosis Report 2022.; 2022.

5. Cassini A, Högberg LD, Plachouras D, Quattrocchi A, Hoxha A, Simonsen GS, et al. Attributable deaths and disability-adjusted life-years caused by infections with antibiotic-resistant bacteria in the EU and the European Economic Area in 2015: a population-level modelling analysis. Lancet Infect Dis (2019) 19(1):56–66. doi:10.1016/S1473-3099(18)30605-4

6. Kuijl C, Savage NDL, Marsman M, Tuin AW, Janssen L, Egan DA, et al. Intracellular bacterial growth is controlled by a kinase network around PKB/AKT1. Nature (2007) <otherinfo>4</otherinfo> 50(7170):725–730. doi:10.1038/nature06345

7. Maiti D, Bhattacharyya A, Basu J. Lipoarabinomannan from <em>Mycobacterium tuberculosis</em> Promotes Macrophage Survival by Phosphorylating Bad through a Phosphatidylinositol 3-Kinase/Akt Pathway *. Journal of Biological Chemistry (2001) 276(1):329–333. doi:10.1074/jbc.M002650200

8. Hao-Chieh C, K. KS, Shilpa S, Dasheng W, S. GJ, S. SL, et al. Eradication of Intracellular Salmonellaenterica Serovar Typhimurium with a Small-Molecule, Host Cell-Directed Agent. Antimicrob Agents Chemother (2009) 53(12):5236–5244. doi:10.1128/AAC.00555-09

9. Korbee CJ, Heemskerk MT, Kocev D, van Strijen E, Rabiee O, Franken KLMC, et al. Combined chemical genetics and data-driven bioinformatics approach identifies receptor tyrosine kinase inhibitors as host-directed antimicrobials. Nat Commun (2018) 9(1):358. doi:10.1038/s41467-017-02777-6

10. Napier RJ, Rafi W, Cheruvu M, Powell KR, Zaunbrecher MA, Bornmann W, et al. Imatinib-Sensitive Tyrosine Kinases Regulate Mycobacterial Pathogenesis and Represent Therapeutic Targets against Tuberculosis. Cell Host Microbe (2011) 10(5):475–485. doi:10.1016/j.chom.2011.09.010

11. Bruns H, Stegelmann F, Fabri M, Döhner K, van Zandbergen G, Wagner M, et al. Abelson Tyrosine Kinase Controls Phagosomal Acidification Required for Killing of Mycobacterium tuberculosis in Human Macrophages. The Journal of Immunology (2012) 189(8):4069–4078. doi:10.4049/jimmunol.1201538

12. Jayaswal S, Kamal MdA, Dua R, Gupta S, Majumdar T, Das G, et al. Identification of Host-Dependent Survival Factors for Intracellular Mycobacterium tuberculosis through an siRNA Screen. PLoS Pathog (2010) 6(4):e1000839.

13. Wong D, Bach H, Sun J, Hmama Z, Av-Gay Y. Mycobacterium tuberculosis protein tyrosine phosphatase (PtpA) excludes host vacuolar-H+–ATPase to inhibit phagosome acidification. Proceedings of the National Academy of Sciences (2011) 108(48):19371–19376. doi:10.1073/pnas.1109201108

14. Giver CR, Shaw PA, Fletcher H, Kaushal D, Pamela G, Omoyege D, et al. IMPACT-TB*: A Phase II Trial Assessing the Capacity of Low Dose Imatinib to Induce Myelopoiesis and Enhance Host Anti-Microbial Immunity Against Tuberculosis. *Imatinib Mesylate per Oral As a Clinical Therapeutic for TB. Blood (2019) 134(Supplement_1):1050. doi:10.1182/blood-2019-130275

15. Elkins JM, Fedele V, Szklarz M, Abdul Azeez KR, Salah E, Mikolajczyk J, et al. Comprehensive characterization of the Published Kinase Inhibitor Set. Nat Biotechnol (2016) 34(1):95–103. doi:10.1038/nbt.3374

16. Drewry DH, Wells CI, Andrews DM, Angell R, Al-Ali H, Axtman AD, et al. Progress towards a public chemogenomic set for protein kinases and a call for contributions. PLoS One (2017) 12(8):e0181585-.

17. Tuin AW. Synthesis of New H89 Analogues. In: Synthetic Studies on Kinase Inihbitors and Cyclic PeptidesflJ: Strategies towards New Antibiotics.; 2008:73-94.

18. Korbee CJ, Heemskerk MT, Walburg KV, van den Nieuwendijk R, van Strijen E, Kuijl C, et al. Novel Host-Directed Chemical Compounds Inhibit Intracellular Bacteria by Targeting PCTAIRE Kinases. In: Host-Directed Therapy for Intracellular Bacterial Infections.; 2019:95–120.

19. Verreck FAW, de Boer T, Langenberg DML, van der Zanden L, Ottenhoff THM. Phenotypic and functional profiling of human proinflammatory type-1 and anti-inflammatory type-2 macrophages in response to microbial antigens and IFN-γ– and CD40L-mediated costimulation. J Leukoc Biol (2006) 79(2):285–293. 10.1189/jlb.0105015

20. Kuijl C, Pilli M, Alahari SK, Janssen H, Khoo P-S, Ervin KE, et al. Rac and Rab GTPases dual effector Nischarin regulates vesicle maturation to facilitate survival of intracellular bacteria. EMBO J (2013) 32(5):713–727. 10.1038/emboj.2013.10

21. Takaki K, Davis JM, Winglee K, Ramakrishnan L. Evaluation of the pathogenesis and treatment of Mycobacterium marinum infection in zebrafish. Nat Protoc (2013) 8(6):1114–1124. doi:10.1038/nprot.2013.068

22. Eid S, Turk S, Volkamer A, Rippmann F, Fulle S. KinMap: a web-based tool for interactive navigation through human kinome data. BMC Bioinformatics (2017) 18(1):16. doi:10.1186/s12859-016-1433-7

23. Vaughan TG. IcyTree: rapid browser-based visualization for phylogenetic trees and networks. Bioinformatics (2017) 33(15):2392–2394. doi:10.1093/bioinformatics/btx155

24. Benard EL, van der Sar AM, Ellett F, Lieschke GJ, Spaink HP, Meijer AH. Infection of zebrafish embryos with intracellular bacterial pathogens. J Vis Exp (2012) (61). doi:10.3791/3781

25. Schindelin J, Arganda-Carreras I, Frise E, Kaynig V, Longair M, Pietzsch T, et al. Fiji: an open-source platform for biological-image analysis. Nat Methods (2012) 9(7):676–682. doi:10.1038/nmeth.2019

26. Lin H, Yamashita DS, Zeng J, Xie R, Wang W, Nidarmarthy S, et al. 2,3,5-Trisubstituted pyridines as selective AKT inhibitors—Part I: Substitution at 2-position of the core pyridine for ROCK1 selectivity. Bioorg Med Chem Lett (2010) 20(2):673–678. 10.1016/j.bmcl.2009.11.064

27. Cronan MR, Tobin DM. Fit for consumption: zebrafish as a model for tuberculosis. Dis Model Mech (2014) 7(7):777–784. doi:10.1242/dmm.016089

28. Meijer A, Spaink H. Host-Pathogen Interactions Made Transparent with the Zebrafish Model. Curr Drug Targets (2011) 12:1000–1017. doi:10.2174/138945011795677809

29. Kilinç G, Saris A, Ottenhoff THM, Haks MC. Host-directed therapy to combat mycobacterial infections*. Immunol Rev (2021) n/a(n/a). 10.1111/imr.12951

30. Haraga A, Ohlson MB, Miller SI. Salmonellae interplay with host cells. Nat Rev Microbiol (2008) 6(1):53–66. doi:10.1038/nrmicro1788

31. Tsao H, Zhang X, Fowlkes K, Haluska FG. Relative Reciprocity of NRAS and PTEN/MMAC1 Alterations in Cutaneous Melanoma Cell Lines1. Cancer Res (2000) 60(7):1800–1804.

32. Wang H, Bi J, Zhang Y, Pan M, Guo Q, Xiao G, et al. Human Kinase IGF1R/IR Inhibitor Linsitinib Controls the In Vitro and Intracellular Growth of Mycobacterium tuberculosis. ACS Infect Dis (2022) 8(10):2019–2027. doi:10.1021/acsinfecdis.2c00278

33. Yuan C, Ai K, Xiang M, Xie C, Zhao M, Wu M, et al. Novel 1-hydroxy phenothiazinium-based derivative protects against bacterial sepsis by inhibiting AAK1-mediated LPS internalization and caspase-11 signaling. Cell Death Dis (2022) 13(8):722. doi:10.1038/s41419-022-05151-7

34. Guarin M, Ny A, De Croze N, Maes J, Léonard M, Annaert P, et al. Pharmacokinetics in Zebrafish Embryos (ZFE) Following Immersion and Intrayolk Administration: A Fluorescence-Based Analysis. Pharmaceuticals (2021) 14(6). doi:10.3390/ph14060576

35. Guarin M, Faelens R, Giusti A, De Croze N, Léonard M, Cabooter D, et al. Spatiotemporal imaging and pharmacokinetics of fluorescent compounds in zebrafish eleuthero-embryos after different routes of administration. Sci Rep (2021) 11(1):12229. doi:10.1038/s41598-021-91612-6

