## Supplementary figures and images for "Identification of kinase inhibitors as potential host-directed therapies for intracellular bacteria"

### Supplementary Figure 1

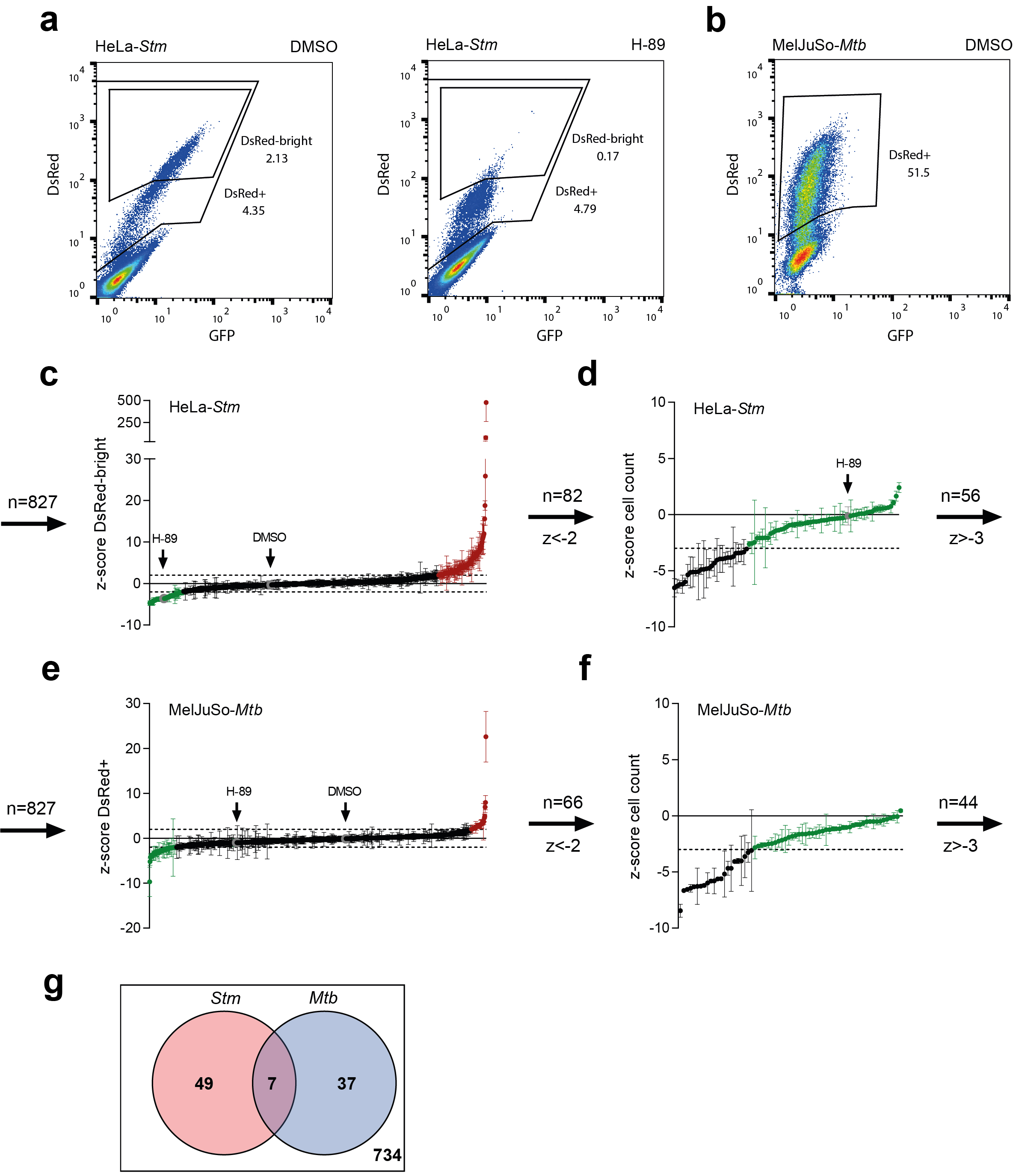

### Supplementary Figure 2

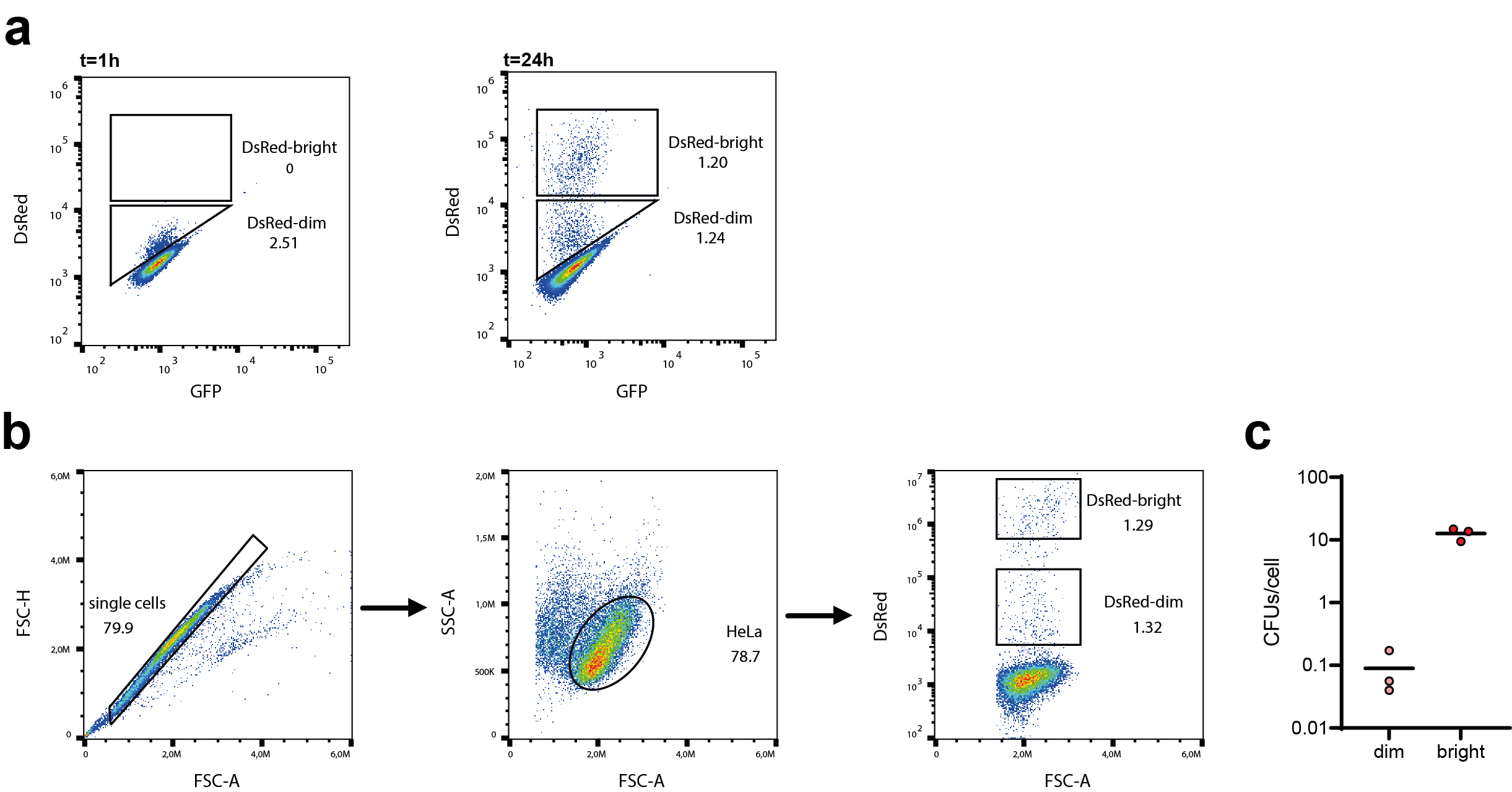

### Supplementary Figure 3

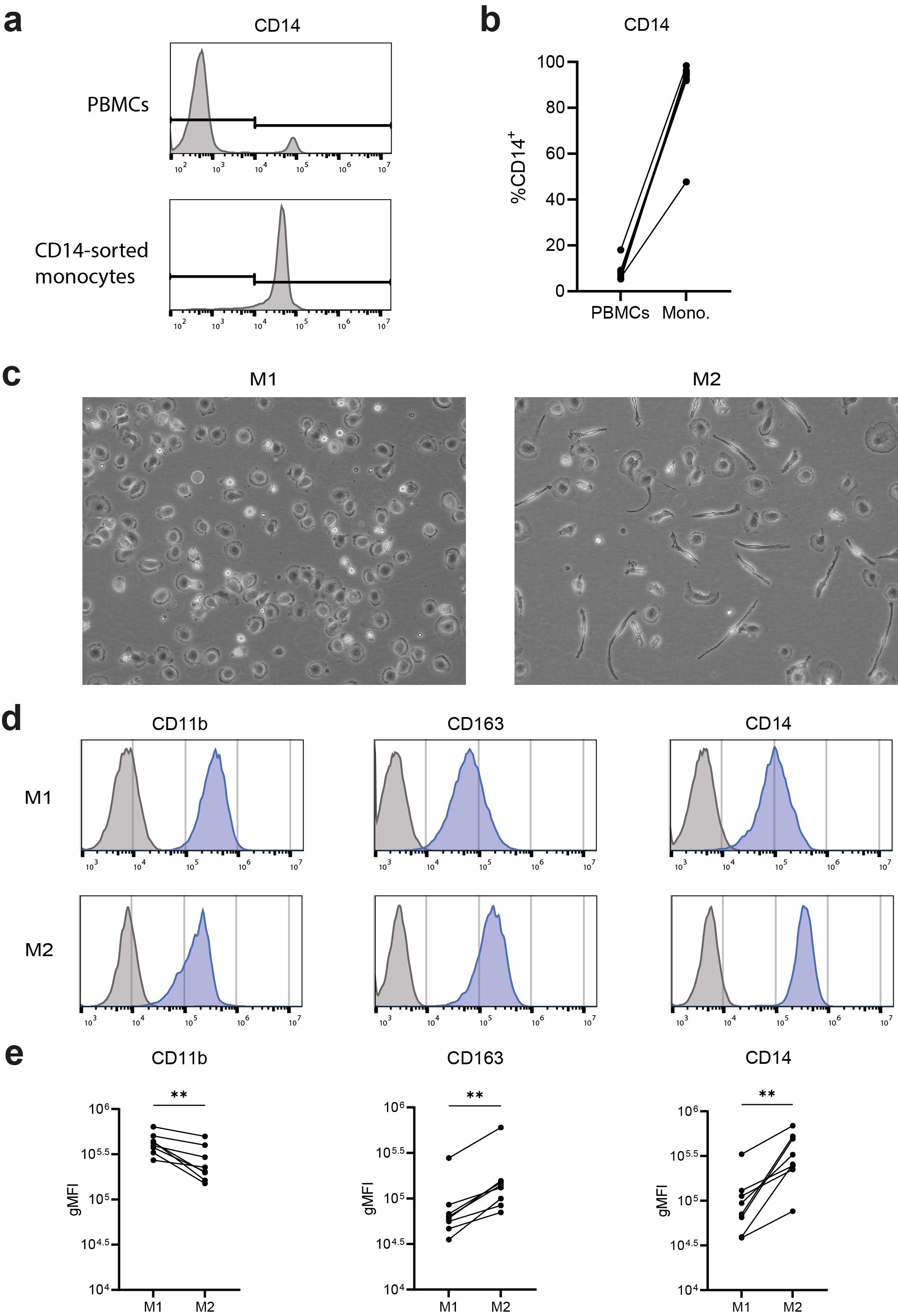
